# Large-scale kinetic metabolic models of *Pseudomonas putida* for a consistent design of metabolic engineering strategies

**DOI:** 10.1101/569152

**Authors:** Milenko Tokic, Ljubisa Miskovic, Vassily Hatzimanikatis

**Affiliations:** Laboratory of Computational Systems Biology (LCSB), EPFL, CH-1015 Lausanne, Switzerland

**Author notes:** **Correspondence:** Ljubisa Miskovic, Laboratory of Computational Systems Biotechnology (LCSB), École Polytechnique Fédérale de Lausanne (EPFL), CH-1015 Lausanne, Switzerland Vassily Hatzimanikatis, Laboratory of Computational Systems Biotechnology (LCSB), École Polytechnique Fédérale de Lausanne (EPFL), CH-1015 Lausanne, Switzerland.

**Keywords:** *Pseudomonas putida*, large-scale and genome-scale kinetic models, nonlinearity, metabolism, thermodynamics, kinetic parameters, uncertainty

## Abstract

A high tolerance of *Pseudomonas putida* to toxic compounds and its ability to grow on a wide variety of substrates makes it a promising candidate for the industrial production of biofuels and biochemicals. Engineering this organism for improved performances and predicting metabolic responses upon genetic perturbations requires reliable descriptions of its metabolism in the form of stoichiometric and kinetic models. In this work, we developed large-scale kinetic models of *P. putida* to predict the metabolic phenotypes and design metabolic engineering interventions for the production of biochemicals. The developed kinetic models contain 775 reactions and 245 metabolites. We started by a gap-filling and thermodynamic curation of iJN1411, the genome-scale model of *P. putida* KT2440. We then applied the redGEM and lumpGEM algorithms to reduce the curated iJN1411 model systematically, and we created three core stoichiometric models of different complexity that describe the central carbon metabolism of *P. putida*. Using the medium complexity core model as a scaffold, we employed the ORACLE framework to generate populations of large-scale kinetic models for two studies. In the first study, the developed kinetic models successfully captured the experimentally observed metabolic responses to several single-gene knockouts of a wild-type strain of *P. putida* KT2440 growing on glucose. In the second study, we used the developed models to propose metabolic engineering interventions for improved robustness of this organism to the stress condition of increased ATP demand. Overall, we demonstrated the potential and predictive capabilities of developed kinetic models that allow for rational design and optimization of recombinant *P. putida* strains for improved production of biofuels and biochemicals.

## 1. Introduction

*Pseudomonas putida* recently emerged as one of the most promising production hosts for a wide range of chemicals, due to its fast growth with a low nutrient [1] and cellular energy [2] demand, considerable metabolic versatility [3], ability to grow in wide range of chemicals [4, 5], suitability for genetic manipulations[6] and its robustness and high flexibility to adapt and counteract different stresses [7]. One of the main advantages of *P. puti*da compared to heavily used industrial workhorses like *E. coli* is its superior tolerance to toxic compounds such as benzene, toluene, ethylbenzene, xylene and other hydrocarbons (e.g., n-hexane and cyclohexane) [8, 9]. For example, Ruhl *at al*. showed that some *P. putida* strains are able to grow in high concentrations of n-butanol [5] up to 6% (vol/vol), whereas the concentrations of 1.5 % (vol/vol) are already toxic for *E. coli* [8].

Recent efforts toward understanding and improving the behavior and systemic properties of *P. putida* metabolism resulted in several genome-scale reconstructions. The first reconstructed Genome-Scale Model (GEM) of *P. putida*, iJN746, was published in 2008 and it comprised 911 metabolites, 950 reactions, and 746 genes [10]. It was rapidly followed by the publication of iJP815 [11] and other reconstructions [12, 13]. The inconsistencies among these models motivated Yuan *et al.* to build so-called pathway-consensus model PpuQY1140 [14]. The so far most complete GEM of *P. Putida*, iJN1411, was published in 2017 by Nogales *et al.* [15], and it contains 2057 metabolites, 2581 reactions, and 1411 genes. These models have been used for studying metabolic features of *P. putida* including the enhanced production of poly-hydroxyalkanoates [16], reconciliation of key biological parameters for growth on glucose under carbon-limited conditions [17], and identification of essential genes for growth on minimal medium [18]. However, stoichiometric models cannot be used to describe the dynamic metabolic responses to changes in cellular and process parameters nor they can consider regulation at the enzyme and post-translational level [19]. We need kinetic metabolic models to address these requirements.

Multiple small-scale kinetic models of *P. putida* metabolism used either of Monod, Haldane and Andrews kinetics for modeling the growth and changes in extracellular concentrations [20–29]. Bandyopadhyay *et al.* used a simple Monod model to study the effect of phenol degradation [22]. Wang and Loh modeled the co-metabolism of phenol and 4-chlorophenol in the presence of sodium glutamate [29], and their kinetic model accounted for cell growth, the toxicity of 4-chlorophenol, and cross-inhibitions among the three substrates. Other models were used for studying growth during benzene [20], toluene [20, 24–26, 28] and phenol biodegradation [20], growth and biosynthesis of medium-chain-length poly-(3-hydroxyalkanoates) [21] and dibenzothiophene desulfurization [23, 27].

More recently, Sudarsan *et al.* developed a kinetic model of the β-ketoadipate pathway that contained mass balance equations for both extracellular and intracellular metabolites described by mechanistic rate expressions based on in vitro investigation of the participating enzymes [30]. Chavarria *et al.* modeled the dynamics of fructose uptake while taking into account the dynamics of gene expression, protein stability, enzymatic activity and the concentrations of intracellular and extracellular metabolites [31].

All these kinetic models are of limited size and with *ad hoc* stoichiometry, which emphasizes a need for developing large-scale kinetic models capable of reliably identifying metabolic engineering targets for production of the desired chemicals [19]. However, construction of large-scale kinetic models remains a challenging task. Each reaction in a kinetic model requires a matching kinetic rate expression along with values of kinetic parameters, which are frequently unknown. Moreover, even if the values of kinetic parameters are available in the literature and databases, their reported values are often spanning several orders of magnitude. Additionally, partial experimental fluxomic and metabolomic data and estimation errors in related thermodynamic properties [19] hinder us from determining unique steady-state metabolic fluxes and metabolite concentrations. As a consequence, we are unable to find a unique model capable of describing the observed physiology. Instead, to overcome this issue, a population of kinetic models is constructed, and statistical methods are used to analyze and predict the metabolic responses in the system [19].

In this work, we first performed a thermodynamic curation and gap-filling of the iJN1411 GEM, and we then systematically reduced this model to derive three different-complexity core models of *P. putida* central carbon metabolism. Next, we applied ORACLE [32–34], a computational framework based on Monte Carlo sampling, to construct large-scale kinetic metabolic models of *P. putida* that were used in two studies: (i) predicting metabolic responses of a wild-type *P. putida* strain to single-gene knockouts; and (ii) improving the responses of this organism to the stress conditions of increased ATP demand. Our results indicate that developed stoichiometric and kinetic models can successfully be used for the design of improved production strains of *P. putida*.

## 2. Results and discussion

### 2.1 Thermodynamically curated genome-scale model of *P. putida*

#### Integration of thermodynamics data

Methods that use thermodynamics data such as the thermodynamics-based flux analysis TFA [35–39] allow us to integrate the metabolomics data together with the fluxomics data, to eliminate in silico designed biosynthetic pathways not obeying the second law of thermodynamics [40, 41], to eliminate infeasible thermodynamic cycles [42–44], and to identify how far reactions operate from thermodynamic equilibrium [45, 46]. Despite the fact that usefulness of thermodynamics has been demonstrated in many applications, only a few reconstructed GEMs are curated for this important property [45, 47–50].

We used Group Contribution method (GCM) [51, 52] to assign the standard Gibbs free energy of formation to 62.3% metabolites and the standard Gibbs free energy of reaction to 59.3% reactions from the iJN1411 model. We calculated the standard Gibbs free energies for all metabolites and reactions participating in the pathways of central carbon metabolism (glycolysis, gluconeogenesis, pentose phosphate pathway, tricarboxylic acid (TCA) cycle). In contrast, we could estimate the standard Gibbs free energy of reaction for only 3.3% reactions in the poly-hydroxyalkanoates (PHA) metabolism because the majority of involved metabolites from these pathways have the structures with unknown residuals which precluded computation of the thermodynamic properties.

#### Integration of physiology data and gap-filling

We integrated experimental measurements of glucose uptake and biomass yield on glucose [53] and metabolite concentrations [54] into the thermodynamically curated model iJN141. The performed TFA indicated that the model predicted ranges of ATP concentrations could not match the values reported in the literature [54, 55]. A reason behind this mismatch could lie in the fact that the H^+^/ATP stoichiometry in the electron transport chain (ETC) of *P. putida* might be inaccurately determined in iJN1411 which would lead to large discrepancies in ATP yield on glucose [3, 56]. Here, we investigated another venue and hypothesized that iJN1411 is missing a critical reaction in the ATP-related metabolism. Therefore, to make model predictions consistent with the experimental observations, we performed gap-filling with the iJO1366 GEM of *E. coli* [57] (Methods). Our analysis indicated that one reaction, sulfate adenyltransferase (SADT2), is missing in the iJN1411. SADT2 plays a role in cysteine formation, and similarly to sulfate adenylyltransferase (SADT), which already exists in the iJN1411, it catalyzes the production of cysteine precursor adenosine 5’-phosphosulfate from ATP and SO4. The production of adenosine 5’-phosphosulfate catalyzed by SADT2 is coupled with GTP consumption, whereas this coupling is absent in SADT. Since the experimental evidence supports that GTP hydrolysis enhances the rate of adenosine 5’-phosphosulfate formation [58], we included this reaction into iJN1411. The thermodynamically curated, gap-filled, model iJN1411 was consistent with the experimental values of both fluxomics and metabolomics data.

### 2.2 Core reduced stoichiometric models of *P. putida*

#### Reconstruction of core reduced models

Using as a basis the curated iJN1411, we employed the redGEM [60] and lumpGEM [61] algorithms to construct a family of three core reduced stoichiometric models of *P. putida* of different complexity. The reduced models were constructed in two steps. First, the redGEM algorithm produced core networks focused around six central carbon subsystems of iJN1411 (glycolysis and gluconeogenesis, pentose phosphate pathway, pyruvate metabolism, TCA cycle and oxidative phosphorylation). Then, the lumpGEM algorithm was used to connect the metabolites of the core networks with 102 biomass building blocks (BBB) of the iJN1411 biomass reaction (Methods).

The simplest out of three core models (subsequently referred to as D1) contained 828 reactions and 286 metabolites distributed over cytosol, periplasm and the extracellular space (Table 1). For 583 out of 828 (70.4%) reactions and 234 out of 286 (81.8%) metabolites from D1 we could calculate the thermodynamic properties (Table 1). The medium complexity core model, D2, contained 704 reactions and 306 metabolites. Out of these, we could calculate the thermodynamic properties for 498 (70.8%) reactions and 253 (82.7%) metabolites. The D3 model had a total of 750 reactions and 336 metabolites with calculated thermodynamic properties for 467 (62.3%) reactions and 276 (82.1%) metabolites (Table 1).

**Table 1.**
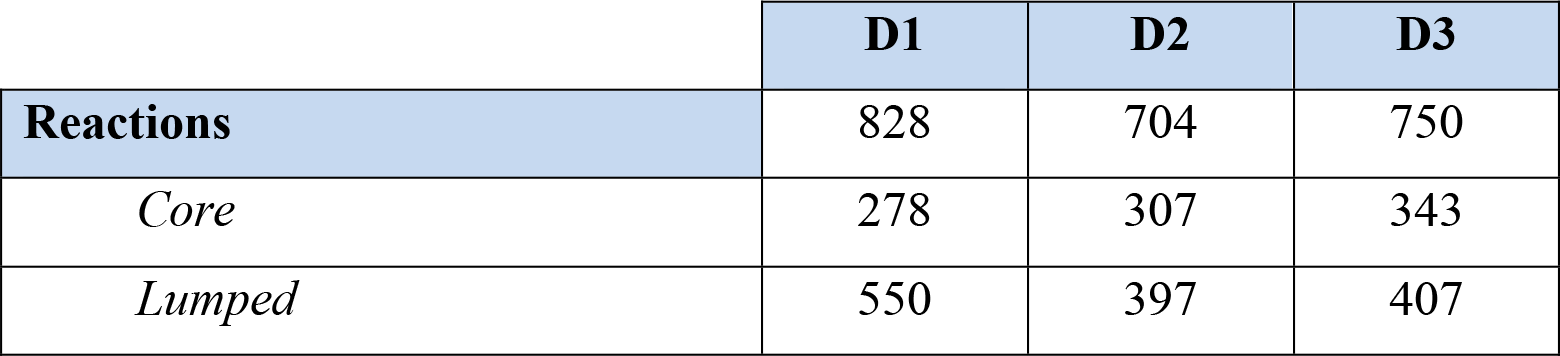
Three reduced core models D1, D2 and D3.

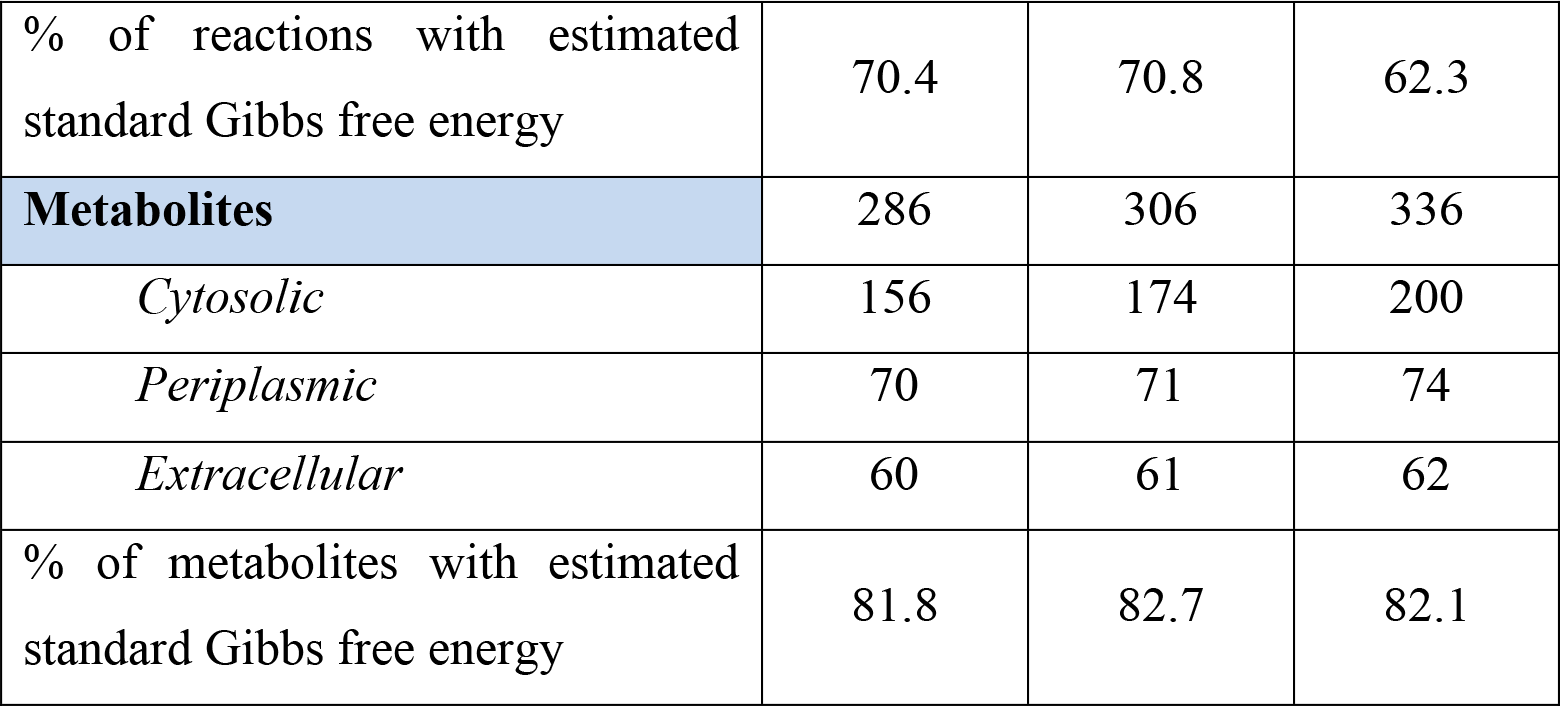

We tested the systemic properties of D1, D2 and D3 against their genome-scale counterpart iJN1441, and we found that they were consistent with the GEM in terms of biomass yields, gene essentiality, and flux and concentration variability (Methods).

#### Essentiality of genes encoding for EDA and EDD

Neither D2 nor the GEM could predict experimentally observed essentiality of genes from the Entner-Doudoroff (ED) pathway. ED pathway is essential for the growth of *P. putida* on glucose, which is experimentally confirmed by the absence of the growth in mutants lacking the key enzymes 2-dehydro-3-deoxy-phosphogluconate aldolase (EDA) and 6-phosphogluconate dehydratase (EDD) [53, 62, 63]. *In silico*, these genes are not essential [18] because the model can replenish the pool of triose phosphates through pentose phosphate pathway.

We analyzed how the directionalities of reactions from the pentose phosphate pathway impact the essentiality of EDA and EDD in D2. We found that the directionalities of three reactions that have glyceraldehyde 3-phosphate (g3p) as reactant (transaldolase, TALA, and two transketolases, TKT1 and TKT2) determine if EDD and EDA are *in silico* essential. When directionality of TKT2 was imposed towards production of g3p, TALA and TKT1 became exclusively unidirectional towards consumption of g3p and production of g3p, respectfully (Fig. 2a), and EDA and EDD were not essential. In contrast, when TKT2 operated towards consumption of g3p EDA and EDD were essential regardless the directionality of the other two reactions (Fig 2b). Therefore, to ensure the consistency of *in silico* and experimentally observed gene essentiality of EDD and EDA in the subsequent studies we imposed the directionality of TKT2 towards consumption of g3p.

**Figure 1.**
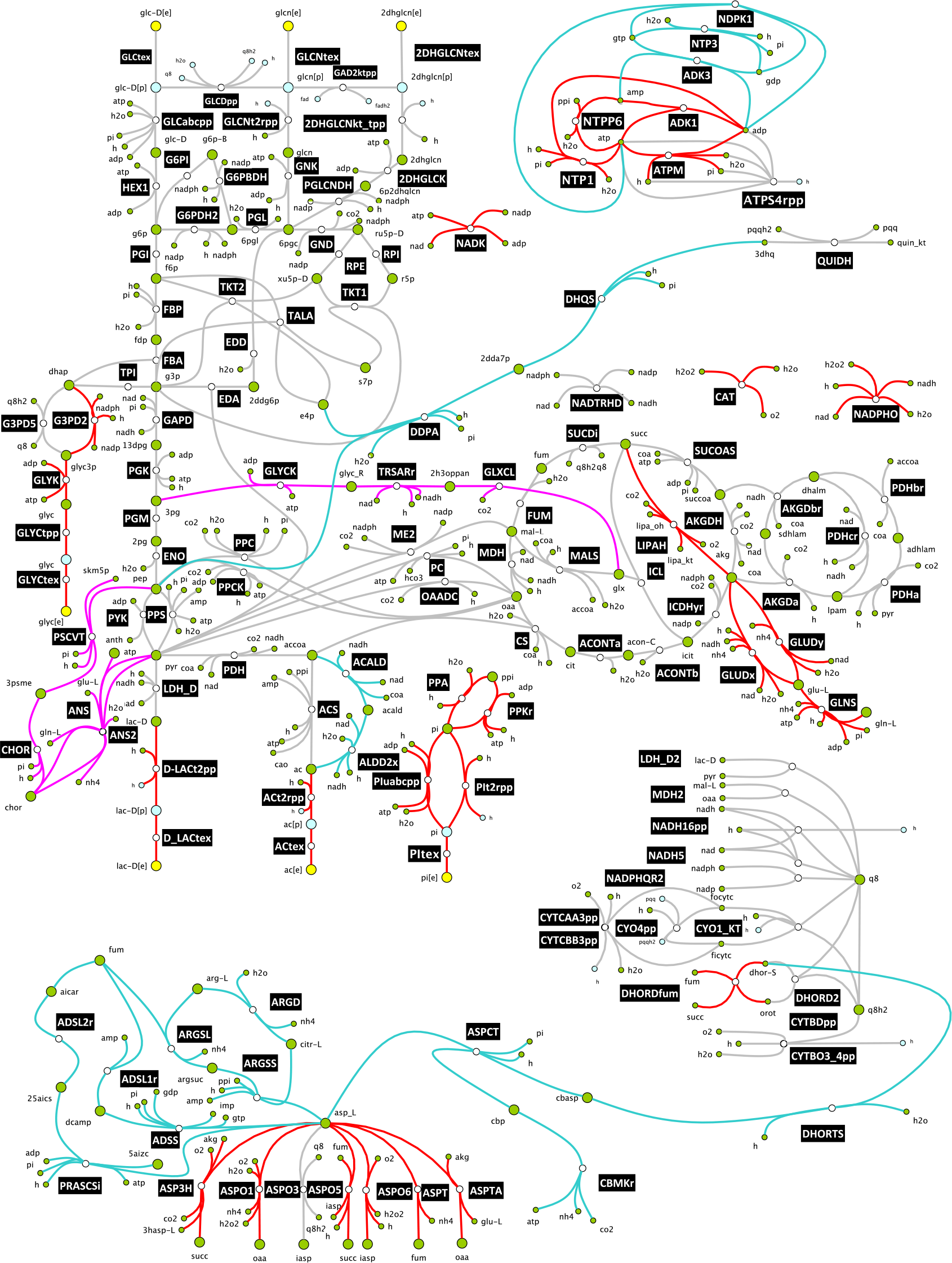
The core networks generated by the redGEM algorithm from iJN1411 genome-scale model. The core network was built around reactions (grey) that belong to the six subsystems of central carbon metabolism (glycolysis and gluconeogenesis, pentose phosphate pathway, pyruvate metabolism, TCA cycle and oxidative phosphorylation). Reactions belonging to one-reaction step, two-reaction-step, and three-reaction-step connections between the six subsystems are marked in red, cyan and magenta, respectively. The stoichiometry of the reduced models and a complete list of reactions and metabolites are provided in Supplementary Files S1-S3.

**Figure 2.**
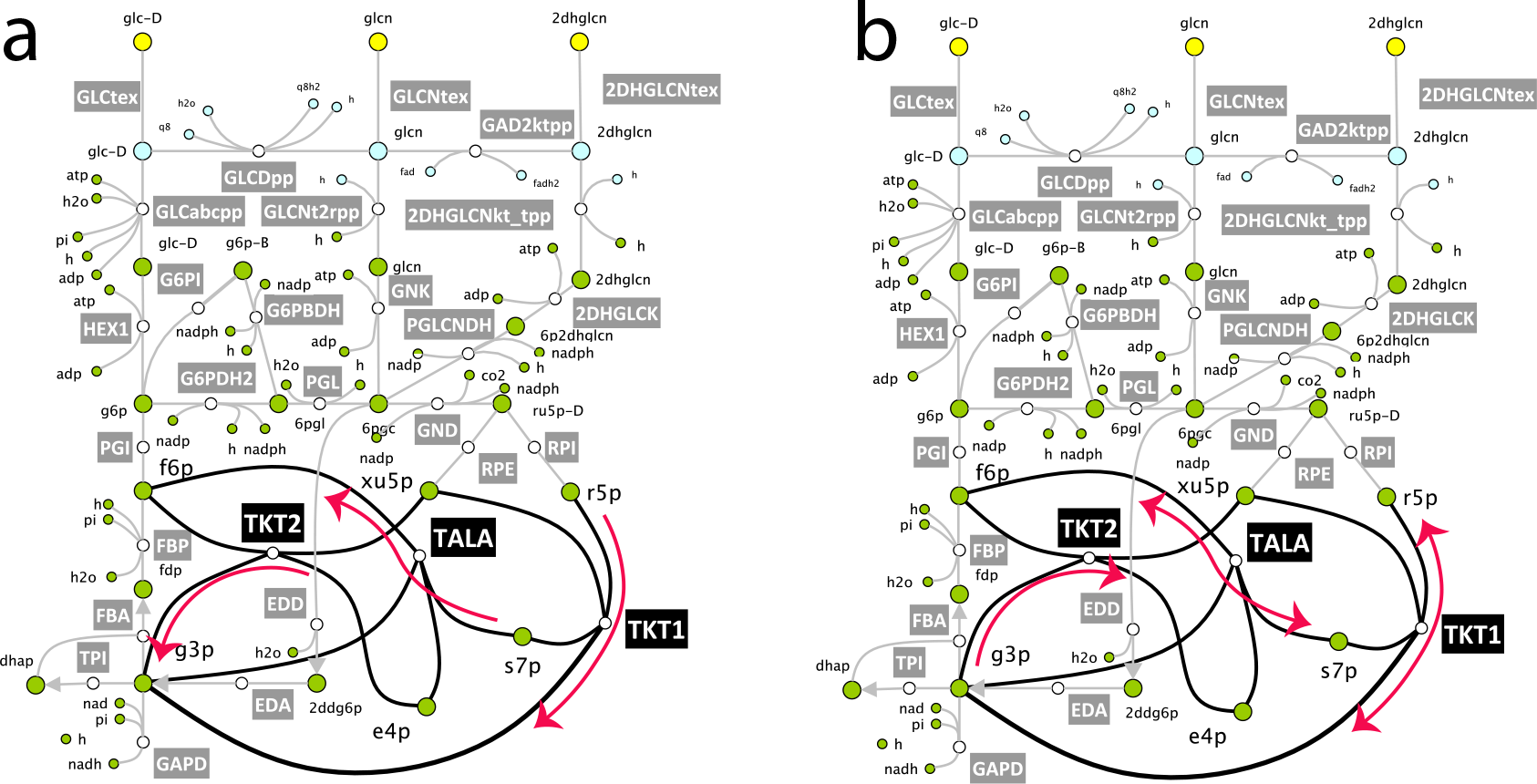
The directionality of transketolase 2 (TKT2) impacts the *in silico* essentiality of two genes encoding EDD and EDA from the Entner-Doudoroff pathway. **(a)** if TKT2 operates towards production of g3p, then due to the stoichiometric coupling transketolase 1 (TKT1) and transaldolase (TALA) are unidirectional and EDD and EDA are not *in silico* essential. **(b)** if TKT2 operates towards consumption of g3p, EDD and EDA are *in silico* essential irrespectively of the directionalities of TKT1 and TALA.

### 2.3 Kinetic study of wild type *P. putida* physiology

#### Model responses to six single-gene knockouts

The reduced D2 model was used as a scaffold for constructing a population of thermodynamically feasible kinetic models. We preconfigured this model for kinetic studies (Methods), and we performed TFA. We found that all reaction directionalities within the obtained thermodynamically feasible steady-state flux and metabolite concentration profile were in agreement with the pre-assigned directionalities from iJN1411[15] (Supplementary Table S1).

In the process of the construction of kinetic models, we removed the mass balances for the extracellular metabolites from the stoichiometry because we consider the concentrations of extracellular metabolites as parameters. The mass balances for water and the corresponding transport reactions were also removed. We then assigned a kinetic mechanism to each of the enzyme catalyzed reactions in the model, and we integrated experimental values for 21 Michaelis constants (K_m_’s) that we found for the *Pseudomonas* genus in the Brenda database [64–67]. Next, we used ORACLE [32–34, 68–71] to construct a population of 50’000 nonlinear kinetic models around the computed steady-state flux and concentration profile (Methods). The resulting structure of kinetic models consisted of 775 enzymatic reactions and 245 mass balances for metabolites distributed over cytosol and periplasm.

As a test for evaluating predictive capabilities of the constructed models, we computed the flux control coefficients [72, 73] of glucose uptake and specific growth rate with respect to six enzymes (glucose dehydrogenase (GLCDpp), hexokinase (HEX1), gluconokinase (GNK), EDA, EDD, and phosphogluconate 2-dehydrogenase (PGLCNDH)), and compared them with the experimentally measured responses of the glucose uptake and specific growth rate to single-gene knockouts of these six enzymes [53].

The computed control coefficients for the glucose uptake and specific growth rate were in a qualitative agreement with the data reported by del Castillo *et al.* [53] (Supplementary Table S2), i.e., a decrease in the enzyme activity of the six enzymes would lead to a decrease in both the glucose uptake and specific growth rate (Fig. 3a and 3b). Nevertheless, a closer inspection of the flux control coefficients of glucose uptake revealed that for four enzymes (GNK, EDD, EDA and PGLCNDH) the error bars were spread around zero values (Fig. 3a). This meant that there was a subpopulation of models with inconsistent predictions with some of the six knockouts. In fact, only 4 999 (~10%) out of 50 000 computed models were able to correctly predict responses to all 6 knockouts reported in del Castillo *et al.* [53] due to the large uncertainty in the assigned values of the kinetic parameters. This type of uncertainty remains one of the major difficulties that limit the predictive strength of kinetic models [19].

**Figure 3.**
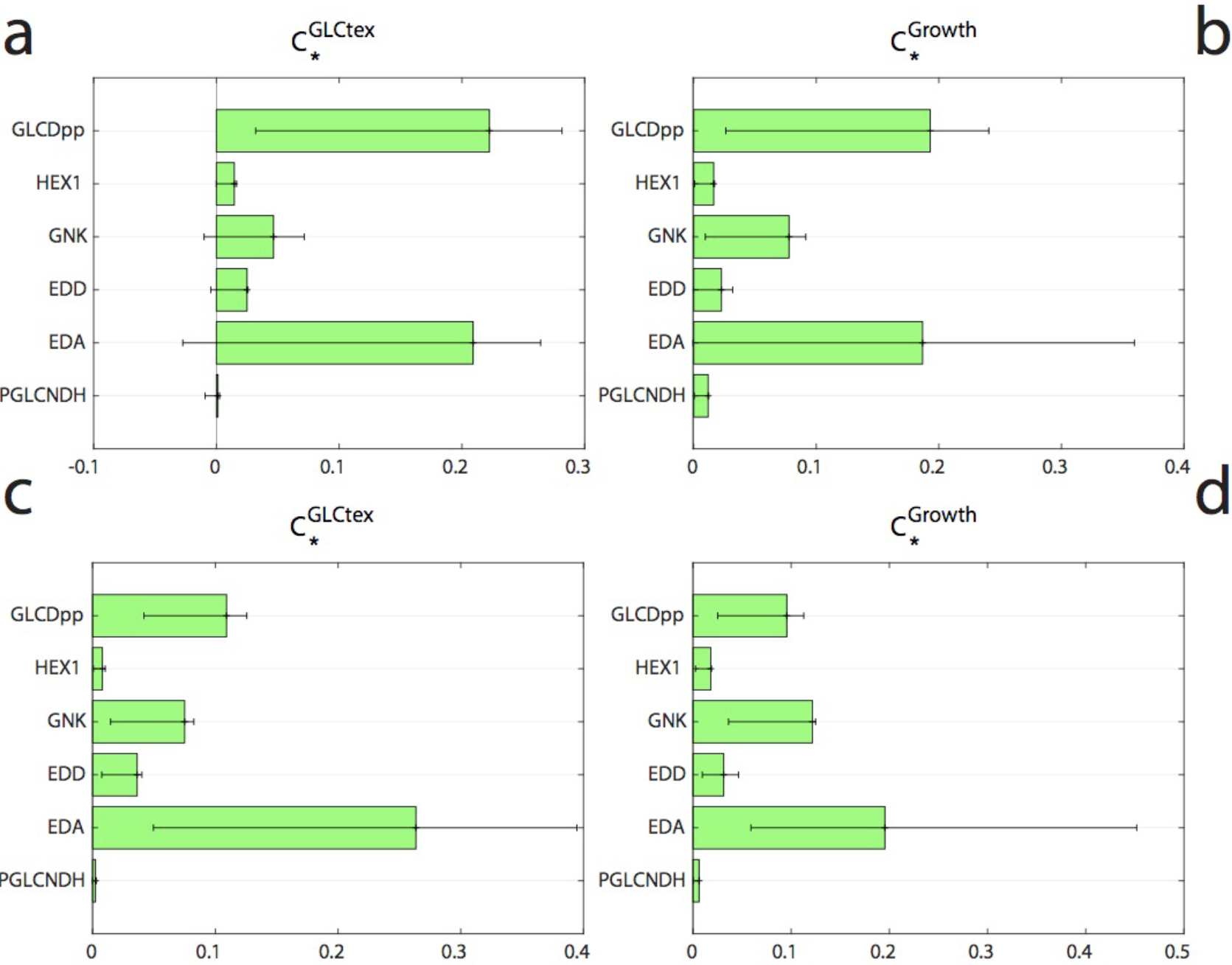
Distribution of the control coefficients of glucose uptake (GLCtex) and specific growth rate (Growth) for the wild-type physiology of *P. putida*. The control coefficients of glucose uptake **(a)** and specific growth rate **(b)** were first computed using an unbiased sampling in ORACLE, and then further refined using the machine learning methodology iSCHRUNK **(c)** and **(d)**. The green bars represent the mean values of the control coefficients, whereas the error bars correspond to the 25 and 75 percentiles of the distributions.

#### Refinement of model responses to six single-gene knockouts

To eliminate the inconsistencies with the experimental data observed for some of the predicted responses, we employed a machine learning method iSCHRUNK [74] (Methods). The method allowed us to identify seven kinetic parameters and their ranges that ensure the consistency of model responses with the experimental observations, and interestingly, all parameters were related with the ED pathway (Table 2).

**Table 2.**
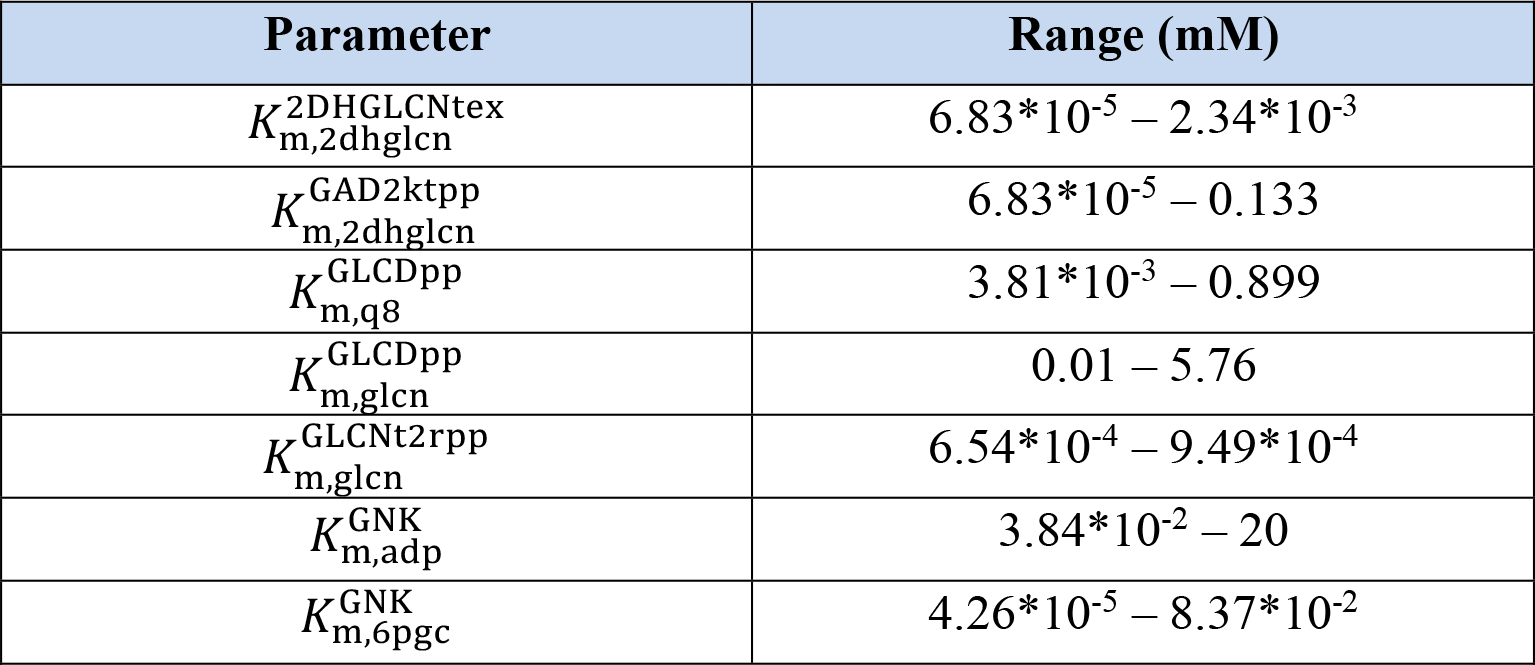
Parameter ranges inferred by the iSCHRUNK method. Abbreviations: 2DHGLCNtex, ketogluconate transport via diffusion extracellular to periplasm, GAD2ktpp, gluconate 2 dehydrogenase periplasm, GLCDpp, glucose dehydrogenase ubiquinone 8 as acceptor periplasm, GLCNt2rpp, D-gluconate transport via proton symport reversible periplasm, GNK, gluconokinase, 2dhglcn, 2-dehydro-D-gluconate, 6pgc, 6-phospho-D-gluconate, adp, ADP, glcn, D-gluconate, q8, ubiquinone-8.

We generated a novel population of kinetic models with ORACLE with constrained ranges of these seven parameters as defined by iSCHRUNK, and we computed the distributions of corresponding control coefficients for the glucose uptake and specific growth rate. Out of 50’000 models, 29’979 (~60%) models correctly predicted the changes in the glucose uptake rate to six single-gene knockouts [53] (Fig. 3c), while 35’955 (~72%) models agreed with the experimental data for the specific growth rate (Fig. 3d). In total, 26’120 (~52%) models were consistent with both the experimental data for the glucose uptake and the specific growth rate.

We discovered with iSCHRUNK that operating regimes of only a few enzymes determine metabolic responses to multiple single-gene knockouts. This emphasizes the significance of accurately determining the kinetic parameters of such important enzymes in order to obtain model responses consistent with the experimental observations. This also implies that we have to consider complex kinetic phenomena such as crowding when modeling kinetic properties of certain enzymes [75].

#### Assessment of estimated kinetic parameters

In the ORACLE framework, we employ the Monte Carlo sampling technique to compute the saturation states of enzymes. We then use these quantities to back-calculate the unknown values of Michaels constants (K_m_’s) [33, 34, 70]. To obtain an unbiased assessment of the accuracy of our estimates, we recomputed 50’000 models without imposing the experimentally available values of K_m_’s from the BRENDA database [64–67]. Comparison of our estimates against available values of K_m_’s from BRENDA showed that ORACLE could capture the ranges for 17 out of 21 K_m_’s (Fig. 4). Considering that in the estimation process we did not use any kinetic parameters values and that the underlying system is undetermined, this result is remarkable because it indicates that ORACLE with integrated fluxomics and metabolomics data together with the physico-chemical laws is capable to provide consistent estimates for a large number of kinetic parameters. This means that ORACLE estimates can be used as hypothetic values for studies where the unknown kinetic parameters are required.

**Figure 4.**
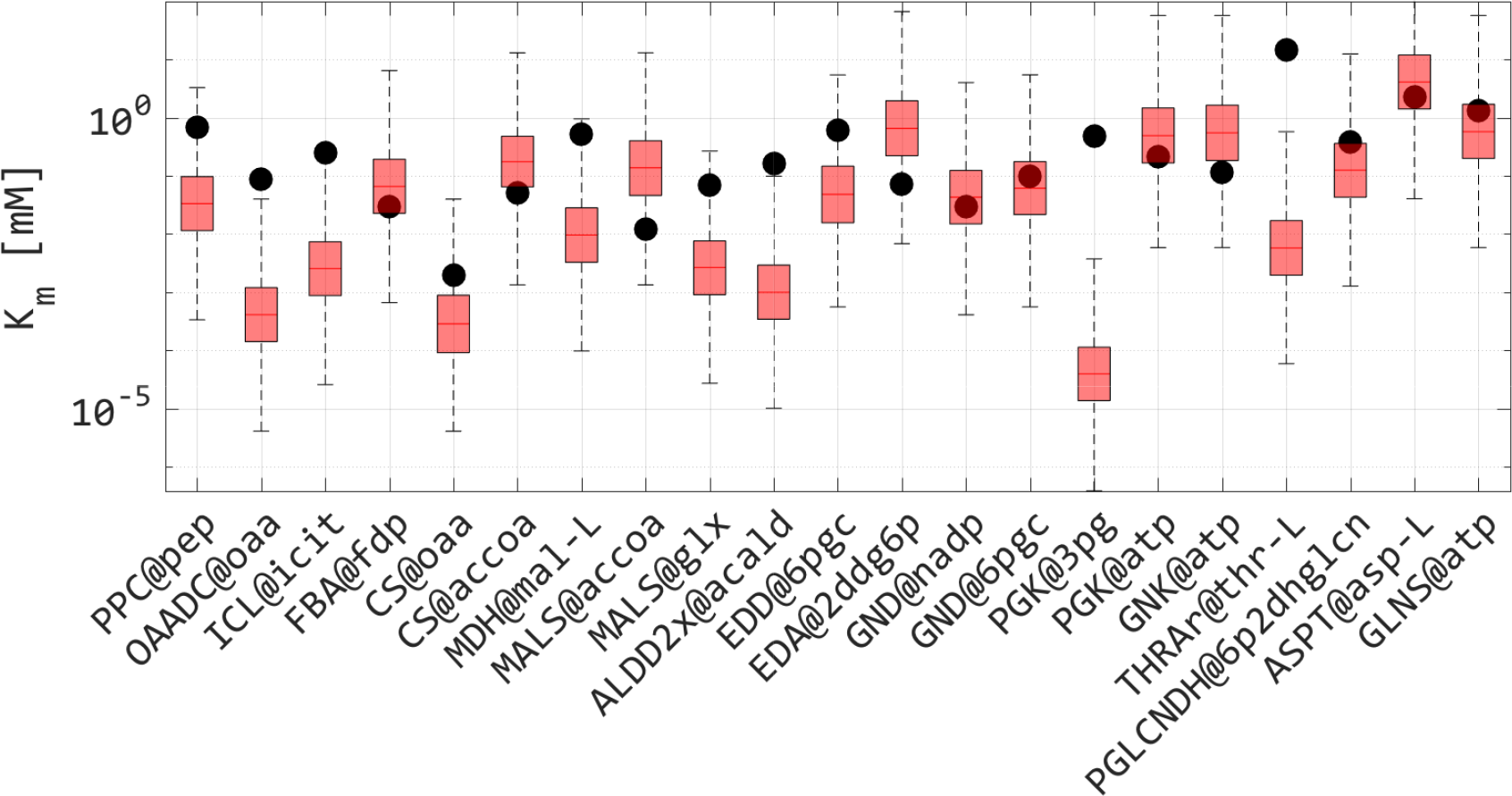
Estimates of Michaelis constants, K_m_’s, predicted by ORACLE. Distribution of K_m_’s estimated with ORACLE (red boxplots) without imposing experimental values from BRENDA (black dots). Whiskers represent minimal and maximal value predicted by ORACLE. Full names of reactions are provided in Supplementary Table S3

For the four remaining parameters such as Michaelis constant of L-Threonine in threonine aldolase or isocitrate in isocitrate lyase, ORACLEs underestimated experimental values up to one and half orders of magnitude (Fig. 4). The discrepancies between the estimated and measured values of these parameters can originate from different sources: (i) the K_m_ values from BRENDA were measured on several different species from the *Pseudomonas* genus, whereas our K_m_ values were estimated using a *P. putida* model and the experimental data were acquired on *P. putida* (fluxomics data) and *P. taiwanensis* (metabolomics data); and (ii) large uncertainty in available and partially available experimental data. In general, the more experimentally measured data are available for integration in the models by ORACLE, the better their predictive capability will be.

### Kinetic study of increased ATP demand in *P. putida*

One of the main advantages of *P. putida* over *E. coli* is its capacity to adjust to different environmental challenges without showing a distinct phenotype [7]. Ebert and co-workers investigated the impact of increased ATP hydrolysis on the *P. putida* metabolism by titration of 2,4-dinitrophenol (DNP), and they demonstrated that DNP concentrations below 300 mg/l did not impact the specific growth rate of *P. putida* [7]. In comparison, *E. coli* shows a significant reduction in the specific growth rate already at the concentrations of 138 mg/l [76]. Above the concentration of 300 mg/l, DNP caused a significant reduction of *P. putida’s* specific growth rate and increase of the glucose uptake (Figure 5a and b). At the concentration of 700 mg/l of DNP, glucose uptake reached the maximum of ~11 mmol/gDCW/h. For larger values of DNP concentration, both the glucose uptake and the specific growth rate declined.

**Figure 5.**
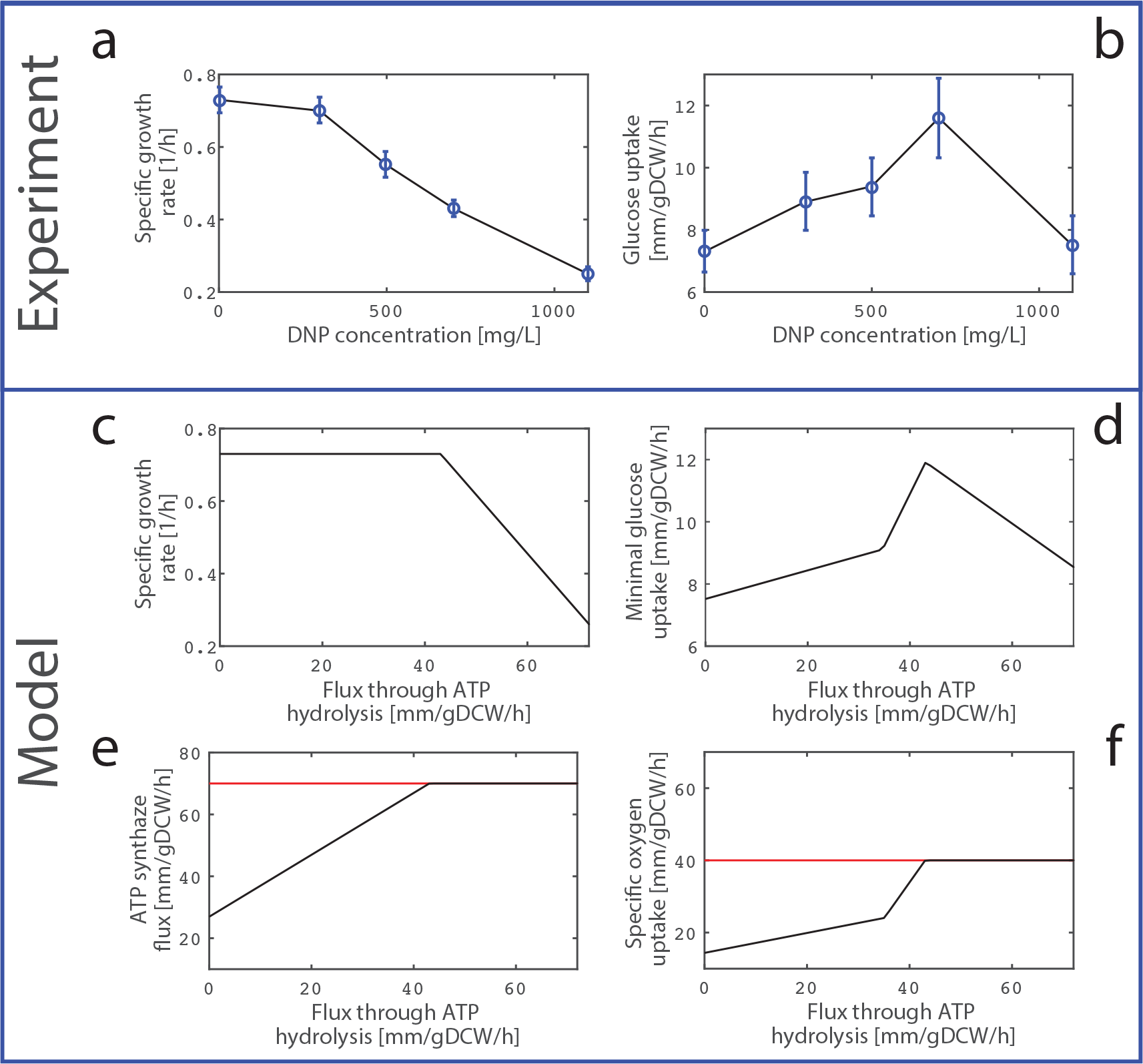
Fermentation profile of *P. putida* metabolism under increased ATP demand. Experimentally measured specific growth rate **(a)** and glucose uptake rate **(b)** of *P. putida* as the ATP demand induced by titration of 2,4 dinitrophenol (DNP) increases. The profiles of specific growth rate **(c)**, glucose uptake rate **(d)**, flux through ATP synthase **(e)** and oxygen uptake rate **(f)** computed by TFA.

#### Modeling and TFA of increased ATP demand

We preconfigured the model for this study (Methods) and used it to simulate the impact of increased ATP demand on the *P. putida* metabolism by gradually increasing the minimally required flux through ATP hydrolysis in increments of 1 mmol/gDCW/h (Fig. 5). We set the upper bound of the specific growth rate to 0.73 1/h, as reported in Ebert *et al.* [7] for the DNP concentration of 0 mg/l. Based on the performed sensitivity analysis of model responses to upper constraints on the oxygen uptake rate and ATP synthase (Methods), we set the upper bounds on the oxygen uptake rate and ATP synthase to 40 mmol/gDCW/h and 70 mmol/gDCW/h, respectively. The glucose uptake rate was left unconstrained.

In agreement with the experiments, the model predicted that the minimal glucose uptake of 7.51 mmol/gDCW/h is required to attain the specific growth rate of 0.73 1/h when the lower bound of the flux through ATP hydrolysis is set to 0 mmol/gDCW/h (Fig. 5c and 5d). Also consistent with the experiments, with the increase of the minimally required ATP hydrolysis flux, the required minimal glucose uptake was increasing (Fig. 5d) simultaneously with an increase of the ATP synthesis flux and minimal oxygen uptake (Fig. 5e and 5f), while the specific growth rate remained stable (Fig. 5c). For the ATP hydrolysis flux of 37 mmol/gDCW/h, the minimal glucose uptake was 9.56 mmol/gDCW/h and the slope of the minimal glucose and oxygen uptake became steeper (Fig. 5d and 5f). When the ATP hydrolysis flux reached 44 mmol/gDCW/h, the oxygen uptake rate and ATP synthase flux simultaneously attained their upper bounds (Fig. 5e and 5f). The corresponding minimal glucose uptake was 11.89 mmol/gDCW/h, which was consistent with Ebert *et al.* [7] (11.6 ± 1.2 mmol/gDCW/h). After this point, the required minimal glucose uptake started to decline (Fig. 5d) together with a decline in the specific growth rate (Fig. 5c). For the ATP hydrolysis flux of 73 mmol/gDCW/h, the model predicted the specific growth rate of 0.25 1/h and the minimal glucose uptake rate of 8.54 mmol/gDCW/h, which was slightly more than what was reported in the Ebert *et al.* [7] (7.5 ± 0.8 mmol/gDCW/h).

The thermodynamically-curated core stoichiometric model described well the qualitative behavior of *P.putida* in the stress condition of increased ATP demand. However, the model failed to capture a decrease of the specific growth rate for DNP concentrations in the range of 300-700 mg/l (Fig. 5c). A possible explanation for this discrepancy is that the decrease of specific growth rate in this region might be due to kinetic effects that cannot be captured by stoichiometric models. It is also important to observe that in Ebert *et al.* [7] the increased ATP demand was indirectly induced by tittering different levels of DNP, whereas we simulated that effect by increasing the ATP hydrolysis flux. Since *P. putida* does not necessarily respond to a linear increase in the DNP levels by linearly increasing the ATP hydrolysis, the exact correspondence of the data points in the graphs obtained through experiments and computational simulation was not expected.

#### Improving the robustness of *P. putida* under stress conditions

We then undertook to devise a metabolic engineering strategy that will allow *P. putida* to maintain the specific growth rate for more severe stress conditions. To this end, we computed the steady-state metabolic flux and metabolite concentration vectors for the ATP hydrolysis flux of 44 mmol/gDCW/h. We then built a population of 50’000 kinetic models around the computed steady-state, and we computed the control coefficients for all fluxes and concentrations in the metabolic network.

Analysis of the control coefficients for the specific growth rate revealed several strategies for maintaining high growth in the presence of stress agent 2,4-dinitrophenol which increases ATP demand (Fig. 6). The major positive control over the specific growth at this stress condition have the key enzymes from the Entner-Doudoroff pathway (EDA, EDD and GNK), e.g., the two-fold increase in activity of EDA would improve the specific growth by more than 50%. This control is tightly connected with the ability of ED pathway to generate additional NADPH, necessary to fuel proton-motive-force-driven efflux pumps, the major mechanism of solvent tolerance in *P. putida* [77] or to reduce stress through antioxidant systems that utilize NADPH [78]. Similarly, our analysis suggests that an increase in the activity of GLCDpp that catalyzes the conversion of glucose to periplasmic gluconate would increase the specific growth, i.e., the two-fold increase in GLCDpp activity would result in improved specific growth by ~40% (Fig. 7). Furthermore, reduced activity of aspartate transaminase (ASPTA) or succinate dehydrogenase (SUCDi) would also increase the specific growth.

**Figure 7.**
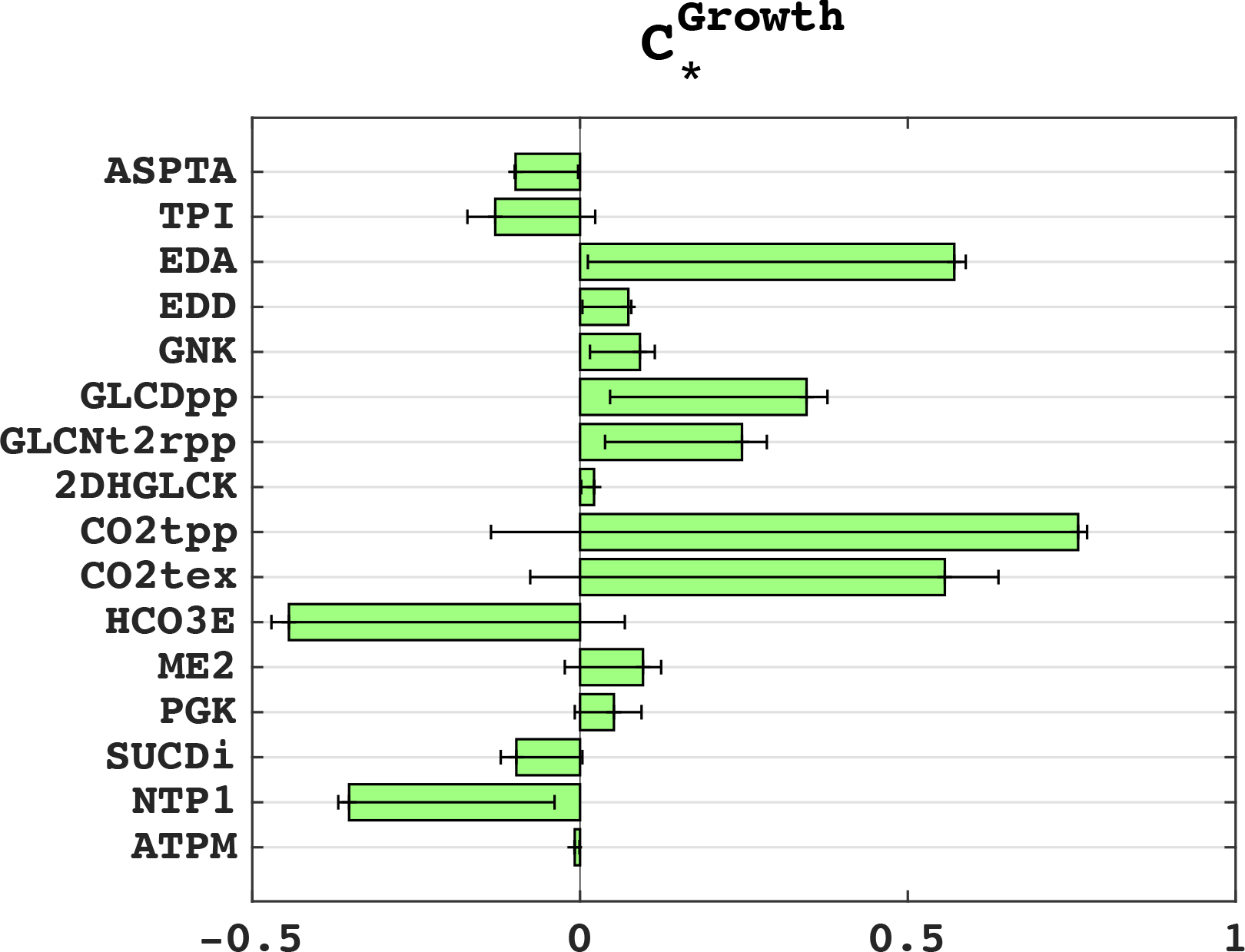
Control coefficients of the specific growth rate in the stress conditions. The green bars are the mean values of the control coefficients, whereas the errorbars correspond to the 25 and 75 percentiles of the distributions.

## 3. Conclusions

This study presents the first thermodynamically curated genome-scale model of *P. putida*. Thermodynamic curation makes the curated GEM iJN1411 amenable for integrating metabolomics data. The integration of thermodynamics data into models restricts the available flux and concentration spaces [35, 39] because thermodynamics determines the directionality in which reactions can operate [35, 37]. For example, Flux Balance Analysis (FBA) performed on iJN1411 indicated that 108 reactions could operate in both forward and reverse direction (bi-directional reactions) while still being consistent with the integrated fluxomics data [53]. However, when additional metabolomics data [54] were integrated with TFA, 21 out of these 108 reactions could not operate in both directions due to thermodynamic constraints (Supplementary Table S4). The thermodynamically curated iJN1411 was further used to develop a family of three systematically reduced models of *P. putida* central carbon metabolism that lend themselves for a wide gamut of metabolic engineering studies.

Current metabolomics measurement techniques do not allow for distinguishing concentrations of the same species in different compartments. Consequently, when integrating metabolomics data in constraint-based techniques that consider thermodynamics such as the energy balance analysis [80], the network-embedded thermodynamic analysis [81] and the thermodynamics-based flux analysis [35, 36, 38, 39, 45], it is commonly assumed that the concentrations of a metabolite appearing in several compartments are identical and constrained within experimentally measured values. We proposed here a novel set of constraints within TFA that enable integration analysis of metabolomics data without imposing this restrictive assumption. In this formulation, we model concentrations of metabolites that exist in several compartments as separate entities, and, at the same time, we preserve the consistency of their values with experimentally measured values for the whole cell. This way, we ensure that the set of possible metabolic outcomes predicted by the model encompasses the actual cellular physiology.

Finally, we derived here the kinetic models of *P. putida‘s* central carbon metabolism containing 775 reactions and 245 metabolites that comprise pathways from glycolysis and gluconeogenesis, pentose phosphate pathway, pyruvate metabolism, TCA cycle, and oxidative phosphorylation. Considering their size, scope, and level of details, the derived models are the largest kinetic model of this organism available in the literature to this date. The potential applications of the developed kinetic models were illustrated in two studies of *P. putida* metabolism.

## Methods

### Integration of metabolomics data while considering cellular compartments

Here we propose a novel set of constraints that allow for concentrations of the same species across different compartments to be different while maintaining the consistency with the experimental measurements.

For the concentration *C*_*M*_ of a metabolite M measured in the range 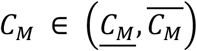 we have:

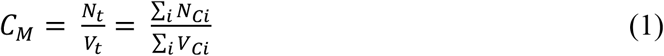

where *N*_*t*_ is the number of moles of M and *V*_*t*_ is the total volume of the cell. *N*_*Ci*_ and *V*_*Ci*_ are the corresponding quantities in compartments *i*. Considering that ∑_*i*_*V*_*Ci*_ = *V*_*t*_, i.e.,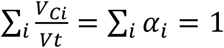, by dividing (1) with *V*_*t*_ we obtain

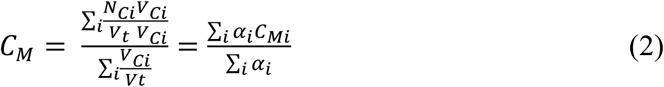

where *C*_*Mi*_ is the concentration of metabolite M in the compartment i and α_i_ is the volume fraction of the compartment i with respect to the entire cell. Observe that α_i_ and *C*_*Mi*_ are positive quantities.

If we apply logarithm to (2), we have:

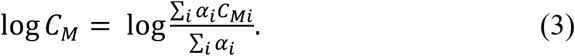

Considering that log is a concave function, we can use Jensen’s inequality [82] where for a concave function *φ* and positive weights *α*_*i*_ it holds that:

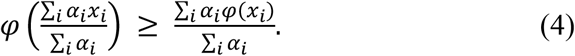

Therefore, by combining (3) and (4) we get:

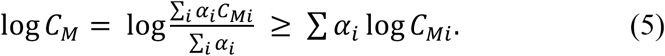

Moreover, if we denote the physiological lower and upper bound on intracellular metabolite concentrations as *LB* = 1 μM and *UB* = 50 mM, respectively, then the upper bound on *C*_*Mi*_, 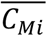, can be derived from the following expression:

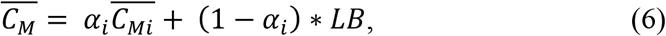

hence

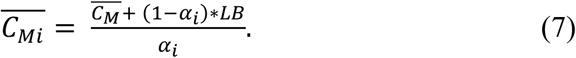

To prevent the case 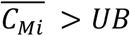 for some values of *α*_*i*_, we put the upper bound on 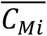 as follows:

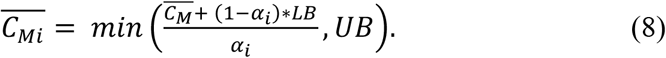

Analogously for the lower bound on the concentration of the metabolite M in the compartment *i*, 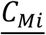, we have:

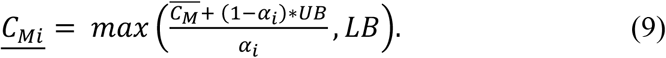

Therefore, instead of using *i* constraints on the compartment species of metabolite M in the form of 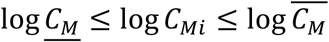, we propose to use *i+2* constraints providing more flexibility and relaxing the assumption on equal concentrations of metabolite M in all compartments:

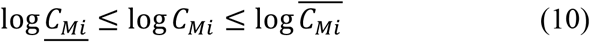

together with (5) and

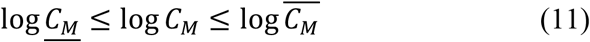

where 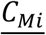 and 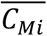 are computed as in (8) and (9).

The volume fractions of cytosol, *α*_1_, and periplasm, *α*_2_, were taken respectively as 0.88 and 0.12 [83].

### Gap-filling of thermodynamically curated iJN1411

We merged two genome-scale models, iJN1411 of *P. putida* and iJO1366 of *E. coli* into a composite model that was used for gap-filling of iJN1411. We removed duplicate reactions from the composite model along with phosphofructokinase (PFK) that is experimentally shown to be absent from *P. putida* metabolism [55]. Compared to iJN1411 the composite model had additional 1201 reactions originating from iJO1366. We imposed experimentally measured ranges of ATP concentrations, glucose uptake and the specific growth rate, and performed TFA while minimizing the number of reactions that can carry flux from the set of the added 1201 reactions. We found from the optimization that it is sufficient to add one out of 1201 reactions (sulfate adenyltransferase (SADT2)) from iJO1366 to iJN1411 to obtain consistency of iJN1411 TFA solutions with the experimental data.

### Systematic reduction of iJN1411

We used the redGEM [60] and lumpGEM [61] algorithms to deliver reduced models of three different sizes (referred in the results section as D1, D2 and D3). The first step in the redGEM algorithm is to select the metabolic subsystems of interest around which the reduced models are built. We selected the following six metabolic subsystems from iJN1411: glycolysis and gluconeogenesis, pentose phosphate pathway, pyruvate metabolism, TCA cycle, and oxidative phosphorylation. From the reactions belonging to these six subsystems, we removed all cofactor pairs and small metabolites such as protons, phosphate groups, and inorganics. We then used a graph search algorithm to identify all one-reaction, two-reaction, and three-reaction steps pairwise connections between six subsystems and formed the core metabolic networks of D1, D2 and D3 model, respectively. We next performed another graph search to find the connections of D1-D3 core networks with the extracellular space. With this step the core networks of D1, D2 and D3 models were finalized.

We then used the lumpGEM [61] algorithm to connect the core networks of D1, D2 and D3 with the building blocks of the iJN1411 biomass reaction. For each of 102 iJN1411 biomass building blocks (BBBs), lumpGEM identified a set of alternative minimal subnetworks that were able to connect precursors belonging to the core network and the BBB. The size of minimal networks is denoted S_min_ [61]. For some studies it is of interest to identify subnetwork of higher sizes. Herein, we identified subnetworks of the size S_min_+2. Finally, lumpGEM collapses the identified subnetworks into lumped reactions, which together with the core networks constitute the core reduced model.

The D1 model consisted of: (i) the D1 core network formed by the reactions and metabolites from the six subsystems and the reactions that belonged to one-reaction-step pairwise connections between these six subsystems [60] (Fig 1); and (ii) lumped reactions that connected the D1 core network with the BBBs. The D2 model contained: the D2 core network containing the D1 core network and the reactions and metabolites that belonged to two-reaction-step pairwise connections between the six subsystems (Fig 1); and (ii) lumped reactions that connected the core network of D2 and the BBBs. The reactions that belonged to the two-reaction-step pairwise connections between the subsystems were predominantly from the fatty-acid and amino-acid metabolism (Supplementary File S2). The core network of the highest complexity model, D3, included also the reactions and metabolites from the three-reaction-step pairwise connections between the six subsystems (Fig 1). The reactions included into the D3 core network were mostly from glyoxylate and dicarboxylate metabolism and folate biosynthesis (Supplementary File S3).

### Consistency checks of core reduced models

We performed a battery of tests to validate the consistency of the systemic properties of the core reduced models D1, D2 and D3 with their GEM counterpart, iJN1411. Here we present and discuss results for D2, the results for D1 and D3 are provided in Supplementary File S4.

We first performed FBA and TFA for the glucose uptake of 10 mmol/gDCW/hr, and we found the identical maximum specific growth rate of μ=0.94 h^−1^ for both D2 and iJN1411, meaning that D2 was able to capture well the physiology of the growth on glucose.

We then carried out the comparison of essential genes between D2 and GEM. *In silico* gene deletion represents one of the most common analysis of metabolic networks, and it is used to assess the predictive potential of the model [10] or to identify main genetic targets for strain engineering [16, 84]. Out of 314 genes that D2 shared with GEM, we identified 47 as *in silico* essential. Out of these 47, 36 were essential in both D2 and GEM and 11 were essential in D2 only (Supplementary Table S5). These 11 genes were essential in D2 because this model was missing some of the alternative pathways from GEM. For example, aceF PP_0338 (encoding for acetyltransferase component of pyruvate dehydrogenase complex) and aceE PP_0339 (encoding for pyruvate dehydrogenase, E1 component) are essential in D2 because they encode for enzymes necessary for synthesizing acetyl-CoA from pyruvate, whereas GEM contains additional alternative pathways for this synthesis. Interestingly, among the 11 genes is tpiA PP_4715 encoding for triose-phosphate isomerase, which is reported as essential in the literature [62].

We next performed thermodynamic-based variability analysis (TVA) on all common reactions and metabolites of D2 and GEM and compared their thermodynamically allowable ranges. We obtained consistent flux ranges for the majority of the reactions, and 131 reactions were less flexible in D2 than in GEM (Supplementary Figure S1). Most of these reactions were in the upper glycolysis such as GAD2ktpp (gluconate 2 dehydrogenase periplasm), GLCDpp (glucose dehydrogenase), HEX 1 (hexokinase) and GNK (gluconokinase), and gluconeogenesis such as PGK (phosphoglycerate kinase), PGM (phosphoglycerate mutase) and ENO (enolase). Additional flexibility of these reactions in GEM comes from the pathways of starch and sucrose metabolism and cell envelope biosynthesis cellulose metabolism, which are absent in D2. The allowable ranges of concentrations of common metabolites of D2 and GEM were consistent. Similar result was reported for the case of *E. coli* where the discrepancy in concentration ranges was reported for only few metabolites [60].

### Preconfiguring stoichiometric model for kinetic studies of wild-type physiology

We expanded the stoichiometric network of D2 by adding the reactions that model free diffusion to extracellular space of all intracellular metabolites that: (i) have less than 10 carbon atoms and do not contain phosphate or CoA; and (ii) do not have an existing transport reaction in the model. This was done to model a possibility that small amounts of these metabolites were produced during fermentation but in insufficient quantities for experimental detection. The expanded model contained 768 reactions and 339 metabolites across cytosol, periplasm, and extracellular space.

Based on the data provided in del Castillo *et al.* [53], we integrated into the model the experimentally measured rates of glucose uptake and biomass growth and we forced the secretion of D-gluconate and 2-dehydro-D-gluconate by putting a lower bound on their exchange reactions to 0.3 mmol/gDCW/hr. For the remaining carbon-based by-products, we allowed only their basal secretion by constraining their transport rates to the extracellular space (10^−6^ - 10^−3^ mmol/gDCW/hr) following the common observation in the literature that *P. putida* can break the carbon down almost without any by-product formation [7]. Furthermore, we integrated 57 experimentally measured intracellular metabolite concentrations [54]. In the model, 12 out of the 57 measured metabolites appear in both cytosol and periplasm. The concentration values of these 12 metabolites were measured per cell and not per compartments, and as discussed previously, to integrate this information for each species in the two compartments only two additional constraints were added in TFA. Overall, these 57 measurements provided constraints for 69 metabolite concentrations in the model.

We then imposed several additional assumptions: (i) TCA cycle was complete [7, 62]; (ii) two glutamate dehydrogenases (GLUDx and GLUDy) were operating towards production of L-glutamate; (iii) dihydrolipoamide S-succinyltransferase was generating NADH from NAD+ [85]; (iv) acetaldehyde dehydrogenase (ACALD) was producing acetaldehyde; (v) ribulose 5-phosphate 3-epimerase (RPE) was converting D-ribulose 5-phosphate to D-xylulose 5-phosphate; (vi) adenylate kinase (ADK1) and nucleoside-diphosphate kinase (NDPK1) were consuming ATP; and (viii) GTP-dependent adenylate kinase (ADK3) was consuming GTP.

### Preconfiguring stoichiometric model for kinetic studies of stress conditions

The stoichiometric model was reconfigured in the following way: (i) we constrained the specific growth rate in the range 0.43 ± 0.2 1/h and the glucose uptake in the range 11.6 ± 1.2 mmol/gDCW/h. These values correspond to the concentration of 700 mg/liter of DNP in the experimental study or 44 mmol/gDCW/h in the simulation study (Figure 5d); (ii) the directionalities of 26 reactions from the glycolysis, gluconeogenesis, PPP and TCA were constrained by putting lower and upper bounds from Ebert *et al.* [7] Interestingly, the reported directionality of TKT2 in this physiological condition was opposite than it was assumed in the study of wild-type physiology; (iii) two glutamate dehydrogenases were operating towards production of L-glutamate; (iv) dihydrolipoamide S-succinyltransferase was operating towards production of NADH from NAD+ [85].

We performed TFA with so configured stoichiometric model, and we found that six reactions (acetaldehyde dehydrogenase acetylating, adenylate kinase, adentylate kinase GTP, sodium proton antiporter, nucleoside diphosphate kinase ATP:GDP and phosphate transport via symport periplasm) could operate in both directions whilst still satisfying the integrated data. To fix the directionalities of these six reactions, we performed another TFA where we minimized the sum of the fluxes in the metabolic network under the constraint that at least 99% of the observed specific growth rate should be attained.

### Sensitivity analysis of metabolic responses to maximal rates in the oxygen uptake and ATP synthesis

Depending on physiological conditions, maximal rates of oxygen uptake and ATP synthase in *P. putida* can take a wide range of values. For instance, in optimally grown *P. putida,* oxygen uptake rate is about 15 mm/gDCW/h [10], while in the stress conditions it can go above 50 mm/gDCW/h [7]. To investigate the effects of the maximal rates on model predictions, we constrained upper bound on biomass growth to 0.73 1/h and we performed multiple TFAs for different combinations of maximal allowed rates of oxygen uptake and ATP synthesis.

We varied the allowed maximal oxygen uptake between 30 and 70 mm/gDCW/h (the range between 40 and 60 mm/gDCW/h was reported in [7]), and the allowed maximal flux through ATP synthase between 40 to 100 mm/gDCW/h. For each combination of oxygen uptake/ATP synthase maximal rates, we computed changes of minimal required glucose uptake with the respect to changes in flux through ATP hydrolysis (Figure 7). For the allowed maximal oxygen uptake of 30 mmol/gDCW/h, the peak of the minimal glucose uptake rate was at 10.22 mmol/gDCW/h, which is slightly under the value reported in Ebert *et al.* [7] (11.6 ± 1.2 mmol/gDCW/h) (Figure 7). For the allowed maximal oxygen uptake of 40 mmol/gDCW/h, the peak of the minimal glucose uptake rate was at 11.89 mmol/gDCW/h which was within the bounds reported in [7], whereas for the allowed maximal oxygen uptake of 50 mmol/gDCW/h, the peak of minimal glucose uptake rate was above the experimental values (13.56 mmol/gDCW/h). Consequently, we used the bound on allowed maximal oxygen uptake rate of 40 mmol/gDCW/h for our kinetic studies.

**Figure 7.**
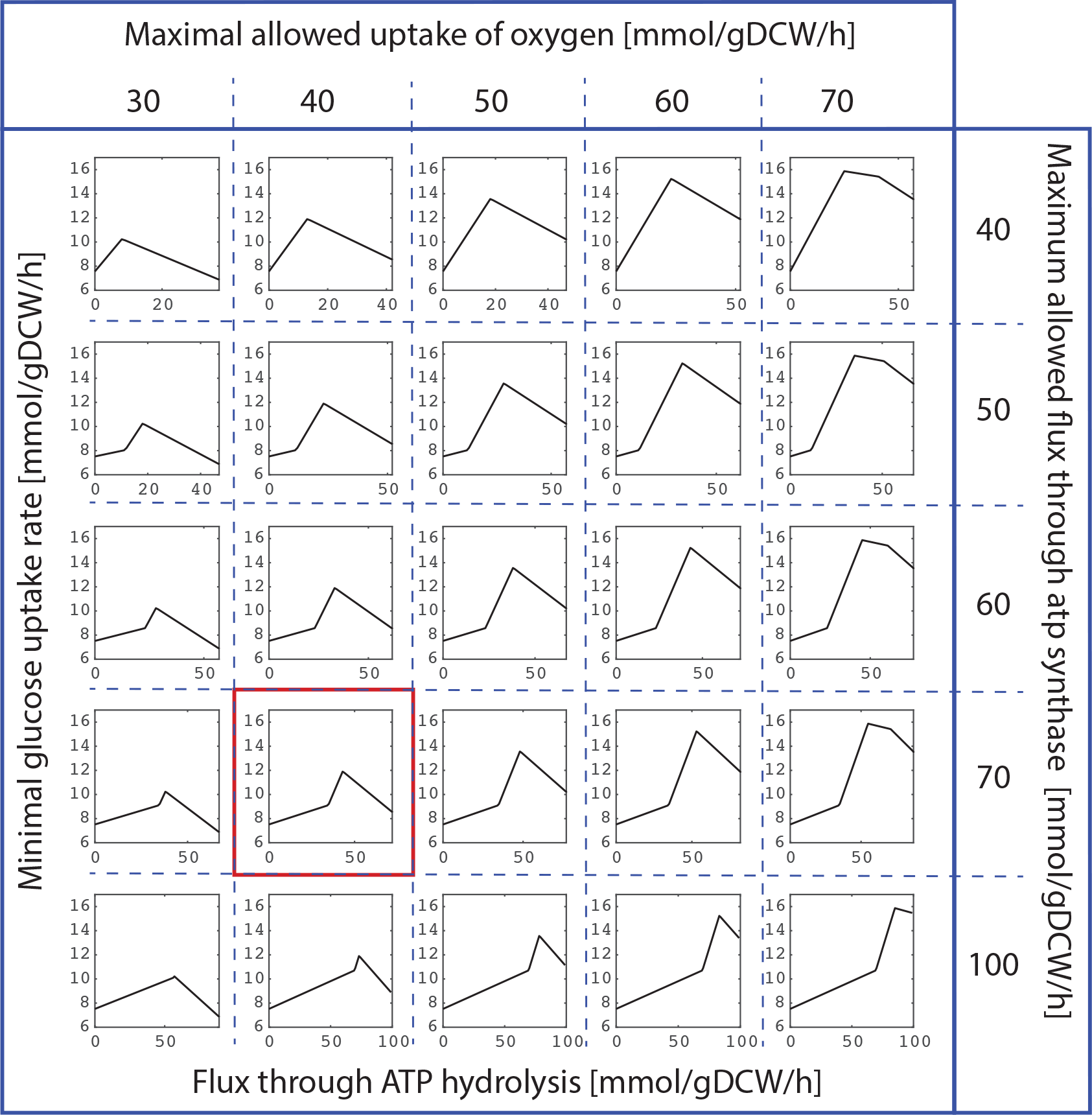
Minimal glucose uptake rate as a function of ATP hydrolysis flux for different combinations of allowed maximal rates of the oxygen uptake and ATP synthesis. The sensitivity analysis indicates that models with the maximal oxygen uptake rate of 40 mmol/gDCW/h and the ATP synthesis rate of 70 mmol/gDCW/h (red box) are providing the best qualitative agreement with the experimental data [7] while maintaining the model flexibility.

Interestingly, the constraint on the allowed maximal ATP synthase rate did not have an effect on the magnitude of the peak value of the minimal glucose uptake rate. Instead, it affected the position of the peak with the respect to the ATP hydrolysis flux (Fig. 7). The higher the ATP synthase rate, the higher ATP hydrolysis flux was required to attain the peak value of the minimal glucose uptake. For example, in the case of the allowed maximal oxygen uptake of 30 mmol/gDCW/h, the ATP hydrolysis flux of 9 and 19 mmol/gDCW/h was required to attain the peak of the minimal glucose uptake of 10.22 mmol/gDCW/h for the allowed maximal ATP synthase rates of 40 and 50 mmol/gDCW/h, respectively. Based on these observations and comparison with the experimental data, one can equally consider values of 50, 60 or 70 mmol/gDCW/h for the upper bound on ATP synthase since all three values describe qualitatively well the experimental data [7] (Fig. 5 and 7). We set the upper bound of ATP synthase to 70 mmol/gDCW/h to keep the maximal flexibility in the model.

## Acknowledgement

M.T. was supported by the ERASYNBIO1-016 SynPath project funded through ERASynBio Initiative for the robust development of Synthetic Biology and the Swiss National Science Foundation grant 315230_163423. L.M. and V.H. were supported by the Ecole Polytechnique Fédérale de Lausanne (EPFL).

## Supporting information

**S1 File: D1 stoichiometric model**

**S2 File: D2 stoichiometric model**

**S3 File: D3 stoichiometric model**

**S4 File: consistency tests for all three reduced models**

**S1 Fig. Flux variability of D2 core carbon subsystems**

**S1 Table: Steady state solution used in building kinetic model**

**S2 Table: Data of 6 single-gene knockouts adapted from del Castillo *et al.[53]***

**S3 Table: Abbreviations for the Figure 4**

**S4 Table: Differences in directionalities between FBA and TFA**

**S5 Table: D2 vs GEM gene essentiality**

## Abbreviations

ORACLE: <u>O</u>ptimization and <u>R</u>isk <u>A</u>nalysis of <u>C</u>omplex <u>L</u>iving <u>E</u>ntities
TFA: <u>T</u>hermodynamics-based <u>F</u>lux <u>B</u>alance <u>A</u>nalysis
GEM: <u>GE</u>nome-scale <u>M</u>odel
MCA: <u>M</u>etabolic <u>C</u>ontrol <u>A</u>nalysis
iSCHRUNK: <u>i</u>n <u>S</u>ilico Approach to <u>CH</u>aracterization and <u>R</u>eduction of <u>UN</u>certainty in the <u>K</u>inetic Models of Genome-scale Metabolic Networks.

