## Supplementary figures and images for "Large-scale kinetic metabolic models of *Pseudomonas putida* for a consistent design of metabolic engineering strategies"

### Supplementary Figure S1

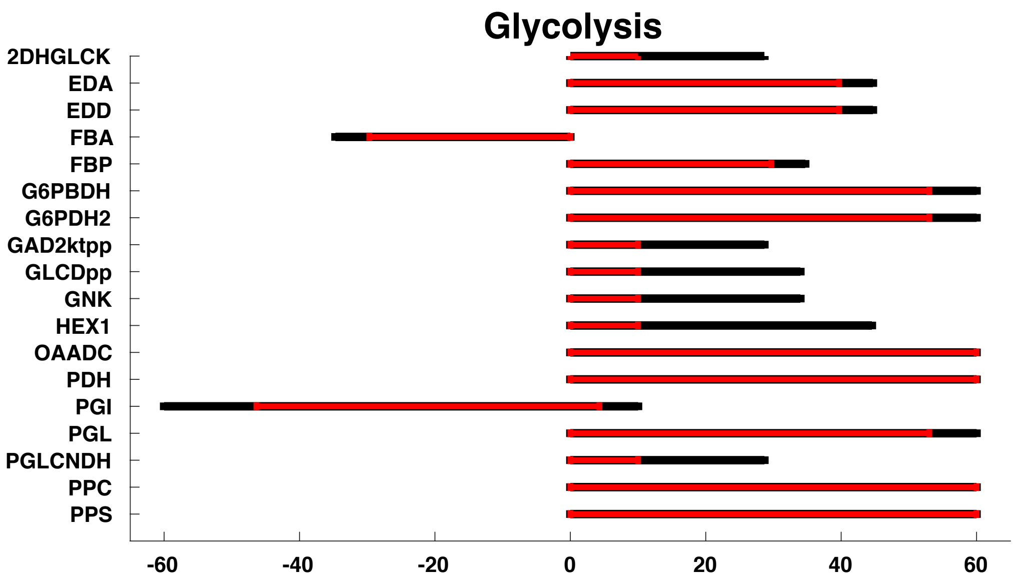

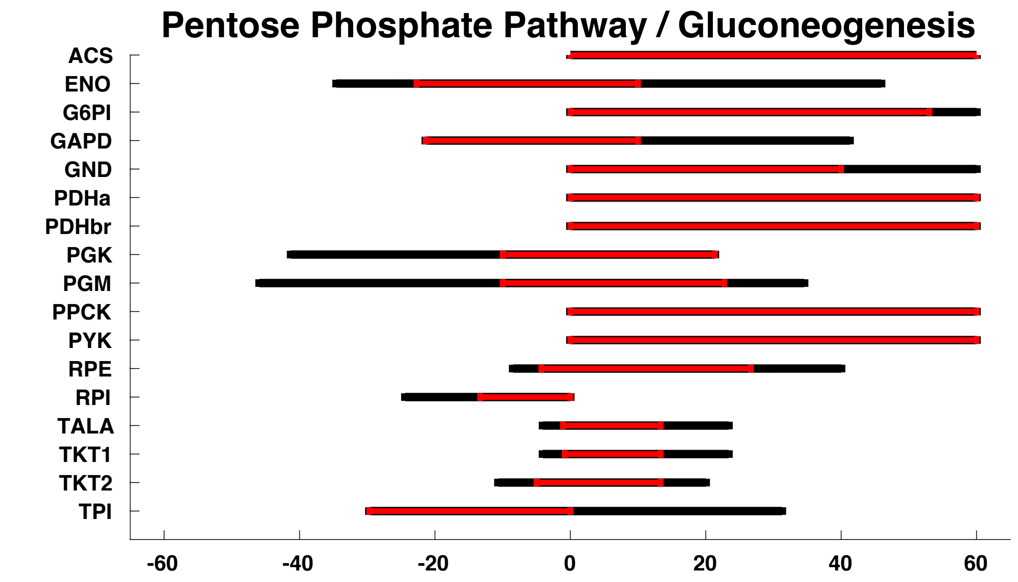

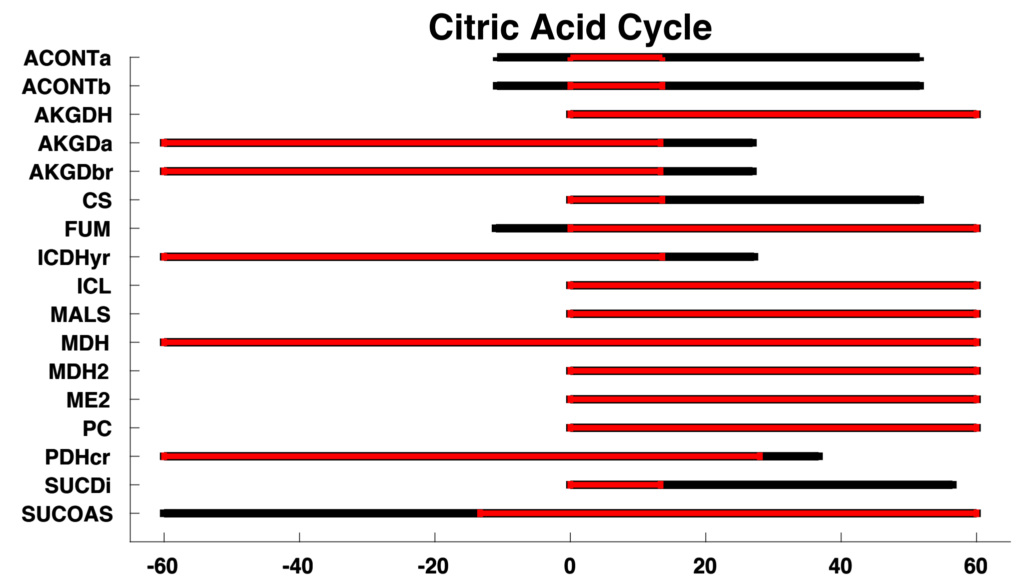
